# Evidence for Multiple Teosinte Hybrid Zones in Central Mexico

**DOI:** 10.1101/2021.02.11.430632

**Authors:** David E. Hufnagel, Kathryn Kananen, Jeffrey C. Glaubitz, José de Jesuś Sánchez-González, John F. Doebley, Matthew B. Hufford

**Author notes:** co-corresponding author: Matthew Hufford, 339A Bessey Hall, Ames, IA. 50011, USA, co-corresponding author: David E. Hufnagel, 1920 Dayton Ave, Bldg 20, Ames, IA. 50010, USA.

## Abstract

- Hybrid zones provide an excellent opportunity for studying population dynamics and whether hybrid genetic architectures are locally adaptive. The genus Zea contains many diverse wild taxa collectively called teosinte. *Zea mays* ssp. *parviglumis*, the lowland progenitor of maize (*Zea mays* ssp. *mays*), and its highland relative *Zea mays* ssp. *mexicana* live parapatrically and, while putative hybrids have been identified in regions of range overlap, these have never been deeply explored.
- Here we use a broadly sampled SNP data set to identify and confirm 112 hybrids between *Zea mays* ssp. *parviglumis* and *Zea mays* ssp. *mexicana*, mostly clustered in three genetically and geographically distinct hybrid groups in Central Mexico.
- These hybrid groups inhabit intermediate environments relative to parental taxa. We demonstrate that these individuals are true hybrids and not products of isolation by distance or ancestral to parviglumis and mexicana. This work expands on previous studies, clearly identifying hybrid zones in *Zea*, genetically characterizing hybrid groups, and showing what appear to be unique genetic architectures of hybridization in distinct hybrid groups.
- With the potential for local adaptation, variable hybrid zone dynamics, and differential architectures of hybridization, we present these teosinte hybrids and parental taxa as a promising model system for studying hybridization and hybrid zones.

## 2 Introduction

Gene flow can homogenize populations across a species range, but can also serve as a major driver of evolutionary change creating novel allele combinations that are then acted upon by selection (Ellstrand, 2014). Hybridization is the creation of genetically distinct individuals through gene flow between differentiated populations; hybrid zones are regions where individuals from these differentiated populations come into secondary contact and reproduce (Barton and Hewitt, 1985). Hybrid zones have been detected in a diverse set of taxa including house mice (*Mus musculus* (Giménez *et al*., 2016)), sunflower (genus *Helianthus* (Rieseberg *et al*., 1999)), Oxford ragwort (genus *Senecio* (Brennan *et al*., 2009)), Heliconius butterflies (genus *Heliconius* (Mallet *et al*., 1990)), and chickadees (genus *Peocile* (Taylor *et al*., 2014)). Genotypes in hybrid zones can be the result of numerous generations of hybridization and recombination and provide an opportunity for evolutionary studies of population dynamics, the timing of secondary contact of parental species, and unequal allele retention including variation in the genetic architecture of hybridization across replicate hybrid zones (Barton and Hewitt, 1985; Smith and O’brien, 2005).

Population dynamics in hybrid zones can vary substantially. For example, hybrid populations can constitute a neutral intergradation zone where hybrids are not at a fitness advantage or disadvantage, a tension zone where hybrids have reduced fitness relative to their parents, or a selection-dependent (i.e., dispersal-independent) zone where hybrids show an increase in fitness relative to parental taxa (Barton and Hewitt, 1985). Due to their deviations from neutrality, tension and selection-dependent zones have received the most theoretical consideration. Factors such as relative migration rate and population density, differential parental fitness and changing climate play a role in determining the size and shape of tension zones (Barton, 1979; Key, 1968; Buggs, 2007; Barton and Hewitt, 1985). A parental population with greater dispersal, population density, or overall fitness tends to push a tension zone away from itself by flooding the zone with its own genotypes (Barton, 1979). In contrast, selection-dependent zones are expected to be wider than tension zones due to increased fitness of hybrids (Barton and Hewitt, 1985) and are also more likely to spread beyond the region where gene flow initiated. Such hybrid populations that become physically isolated may ultimately speciate (Barton and Hewitt, 1985; Abbott *et al*., 2013). For example, *Pinus densata* (High Mountain Pine), *Senecio squalidus* (Oxford Ragwort), and three species resulting from *Helianthus annus* x *H. petiolaris* (*H. anomalus, H. deserticola*, & *H. paradoxus*) were all born out of hybrid zones and subsequently occupied distinct habitats relative to parental populations (Abbott and Brennan, 2014; Abbott *et al*., 2008; Rieseberg *et al*., 1998; Heiser *et al*., 1969).

Additionally, Replicate hybrid zones can be viewed as pools of novel allele combinations upon which selection can act. Patterns of parental allele loss and retention in hybrid populations can be characterized as genomic architectures of hybridization. Nonrandom architectures across hybrid individuals and populations provide evidence that a hybrid zone is selection dominated (Barton and Hewitt, 1985). In the case where multiple hybrid zones are created by similar parental populations and distributed across an environmental gradient, different genomic architectures of hybridization have the potential to be locally adaptive.

Here we establish teosinte (*i*.*e*., wild maize) as a promising system for the study of evolutionary dynamics in hybrid zones and, in particular, adaptive architectures of hybridization. The parental taxa of teosinte hybrids are *Zea mays* ssp. *parviglumis* (hereafter parviglumis) and *Zea mays* ssp. *mexicana* (hereafter mexicana). Parviglumis, best known as the progenitor of domesticated maize (*Zea mays* ssp. *mays*; (Matsuoka *et al*., 2002)), is found in the lowlands of southwest Mexico (*<*1800m), and mexicana is distributed throughout the highlands of the Mexican Central Plateau (1600-2700m; (Hufford *et al*., 2012)). The two subspecies diverged approximately 60,000 generations ago (Ross-Ibarra *et al*., 2009; Hanson *et al*., 1996) and differ in morphological features that suggest local adaptation. Both subspecies also have high diversity and effective population size (Ross-Ibarra *et al*., 2009). Parviglumis plants are green and glabrous, while mexicana individuals are more deeply pigmented and hairy. Differences in pigment and pilosity between parviglumis and mexicana are thought to be associated with adaptation across the altitudinal gradient in western Mexico (Wilkes, 1967; Doebley, 1984). Pigmentation has previously been linked to cold tolerance in maize (Chong and Brawn, 1969; Doebley, 1984) and can be beneficial to high-altitude plants by improving their absorption of radiant solar energy (Galinat, 1967; Chong and Brawn, 1969). Trichome abundance has been associated with plants in cold climates generally (Daubenmire, 1947; Bosabalidis and Sawidis, 2014; Carlquist, 1974). In teosinte, Lauter et al. speculated that macrohairs indurated with silica may form a boundary layer that reduces the loss of absorbed radiant heat at high altitude (Lauter, 2001; Lauter *et al*., 2004).

Putative hybrids between these taxa have been reported at intermediate altitudes in regions of overlap in the distributions of parviglumis and mexicana (Fukunaga *et al*., 2005; van Heerwaarden *et al*., 2011; Pyhajarvi *et al*., 2013). The location of hybrids at intermediate altitudes is compelling since hybrid zones defined by altitude are often adaptive given the substantial environmental variation spanning altitudinal gradients (*e*.*g*., temperature, atmospheric pressure, soil moisture, light intensity, Phosphorous content, and wind velocity (Korner, 2007; Aguirre-Liguori *et al*., 2019)). Moreover, teosinte hybrids have been documented across a region spanning hundreds of kilometers and several degrees of latitude in Mexico, which presents the opportunity for varying adaptive architectures of hybridization. To explore this system more fully and assess the evidence for multiple, independent, hybrid zones, 983 SNPs genotyped in 2,793 individuals from a publicly available data set were analyzed (Fang *et al*., 2012). Additionally, we generated a new data set with a subset of 828 SNPs using the same genotyping platform applied to 239 individuals from 12 populations. In total, we identify a set of 112 parviglumis-mexicana hybrids, residing in three allopatric, genetically distinct hybrid groups in the Central Plateau, the Central Balsas River Valley, and South Guerrero State of Mexico. The environments of these hybrid groups were all shown to be distinct and generally intermediate between that of parviglumis and mexicana. We present multiple, independent sources of evidence that these plants are true hybrids and neither ancestral to parviglumis and mexicana, nor products of Isolation By Distance (IBD). Together, parviglumis and mexicana are a promising system for the study of hybridization and hybrid zones because of their potential for local adaptation within independent hybrid regions, which is potentially linked to differential genome-wide architectures of hybridization.

## 3 Materials and Methods

### 3.1 Acquisition and Generation of SNP Data Sets

Our starting data set was a publicly available collection of 983 SNPs genotyped in 2,793 individuals (van Heerwaarden *et al*., 2011, 2011; Fang *et al*., 2012). From this extensive data set we subsampled parviglumis, mexicana, and also Mexican maize individuals which are interfertile with teosinte and may also represent a source of gene flow. The data set was also filtered by first removing markers, then individuals with *≥* 10% missing data. After data filtering, 967 SNPs genotyped in 1,344 individuals were retained. For retained teosinte individuals, we used the package “dismo” (version 1.1-1) and BIOCLIM data (Hijmans *et al*., 2005) at 30-second resolution to extract environmental data based on geographic coordinates. The same SNP array was used to genotype an additional data set published here. The procedure used to generate these SNPs is described in Weber *et al*. (2007) and van Heerwaarden *et al*. (2010) and the genotyping platform was generated using a discovery panel of 16 teosinte and 14 maize inbred individuals (Wright *et al*., 2005; Weber *et al*., 2007). In the new data set, 828 SNPs were genotyped in 239 individuals. After removing monomorphic loci and applying the same missing data filter used in the 967-SNP data set we retained 820 SNPs in 232 individuals.

### 3.2 Hybrid Identification

For the published 967-SNP data set (Fang *et al*., 2012), we determined whether an individual is substantially admixed by first running a STRUCTURE (version 2.3.4)(Pritchard *et al*., 2000) analysis using group size (k) values from two to six, with 5,000 iterations of burn-in and 10,000 MCMC repetitions. The curve of the likelihood associated with increases in k-values plateaued before reaching k=6. The web application “Structure Harvester” (version 0.6.93) (Earl and vonHoldt, 2011) was used to determine the optimum k (k=3) based on the deltaK statistic, which corresponds to the second order rate of change of the likelihood. Hybrids were defined as individuals with significant (25-50%) attribution to the alternative teosinte subspecies. Teosinte with *≥* 90% self-attribution were considered high-confidence, non-admixed individuals. One plant annotated as parviglumis upon collection showed primarily mexicana ancestry and was removed from the data set.

The published data set is broadly distributed across Mexico but does not include an abundance of individuals sampled per population. Therefore, to evaluate the consistency of hybridization signatures at the population level, an additional STRUCTURE analysis was performed using the more deeply sampled 820-SNP data set. k-values varied from two to eight with 100,000 iterations of burn-in and 1,000,000 MCMC iterations. Higher MCMC iterations were necessary in this analysis to reach convergence. Three was chosen as the most supported k-value for both data sets. Plots were generated using the program distruct (version 1.1) (Rosenberg *et al*., 2002).

### 3.3 Principal Component Analysis

A Principal Component Analysis (PCA) was completed using the package “prcomp” (from the “stats” package version 3.2.2). Monomorphic loci were removed, and data were fit to a standard normal distribution, where missing data has a value of zero, before running “prcomp” (modified from van Heerwaarden *et al*. (2011)).

### 3.4 conStruct Analysis

The program conStruct (version beta) was used to generate a STRUCTURE-like q-matrix while explicitly controlling for isolation by distance. Because the package is not designed for high numbers of individuals and cannot run with more individuals than SNPs, we excluded maize and randomly subsampled one individual from each parviglumis, mexicana, and hybrid population reducing the data set to 195 individuals. The program was run under the spatial model using default parameters at group size two with five replicates. The output from the highest likelihood replicate was then used to determine the percent attribution to the alternative teosinte subspecies for each group.

### 3.5 *F*_*ST*_ x *F*_*ST*_ Calculations and Inversion Analysis

Wright’s *F*_*ST*_ was determined using the R package “hierfstat” (version 0.04-10) for all possible comparisons between high confidence members of the three hybrid groups and high confidence members of the major races of parviglumis (Balsas and Jalisco) and mexicana (Chalco and Central Plateau). Global *F*_*ST*_ values were calculated by averaging all per-locus *F*_*ST*_ values and considering negative values to be zero. Chromosome-level patterns of *F*_*ST*_ for each hybrid group were then compared to parental populations.

Wright’s *F*_*ST*_ was also calculated comparing all high-confidence parviglumis and mexicana samples to each other. Based on chromosomal coordinates and extremely high *F*_*ST*_ values (*>*0.95), four markers were identified as falling within a well known chromosomal inversion on chromosome 4, *inv4m* that distinguishes parviglumis from mexicana ((Pyhajarvi *et al*., 2013); PZD00030.4, PZA01637.3, PZA01637.4, and PZD00030.1). These loci were examined in all high-confidence parviglumis and mexicana samples and subspecies-specific haplotypes were clearly identified. Each individual was then assigned one of five possible inversion states based on genotypes at these loci: P for parviglumis type, M for mexicana type, H for heterozygous type, R for recombinant type, and N for predominantly missing data. P and M were assigned when all loci were either homozygous parviglumis type or homozygous mexicana type, respectively, with one or fewer missing data loci. H was assigned to individuals that were heterozygous at all loci with one or fewer missing data loci. R was assigned to individuals that had a mixture of homozygous parviglumis type loci, mexicana type loci, and heterozygous loci with one or fewer missing data loci, indicating either recombination or sequencing errors. N was assigned to individuals that had two or more missing data loci.

### 3.6 Data Availability

The newly generated 828 SNP data set and information on associated plants and populations can be found at: https://figshare.com/account/home#/projects/98201

### 3.7 Code Availability

All data and scripts needed to replicate our results can be found at: https://github.com/HuffordLab/Hufnagel et al. Teosinte Hybrid Zones

## 4 Results

### 4.1 Identifying Hybrids and Hybrid Groups

To identify hybrids, we used the program STRUCTURE with a subset of a publicly available SNP data set (van Heerwaarden *et al*., 2010, 2011; Fang *et al*., 2012) including 967 SNPs genotyped in 1,344 individuals. STRUCTURE uses a model-based approach to infer population structure and assign individuals to populations probabilistically (Pritchard *et al*., 2000). Results suggested three clear ancestry groups corresponding to parviglumis, mexicana, and maize, with population structure largely consistent across k-values. Based on STRUCTURE results we identified 112 high-confidence hybrids; 634 high-confidence, non-admixed parviglumis individuals; 95 high-confidence, non-admixed mexicana individuals; and 176 parviglumis and 51 mexicana individuals with evidence of low levels of admixture (Fig. 1). Many of the hybrid individuals in both teosinte subspecies were those at mid-elevation and are therefore more likely to have come into contact with and potentially admixed with the other subspecies (Figs 2,S1).

**Figure 1.**
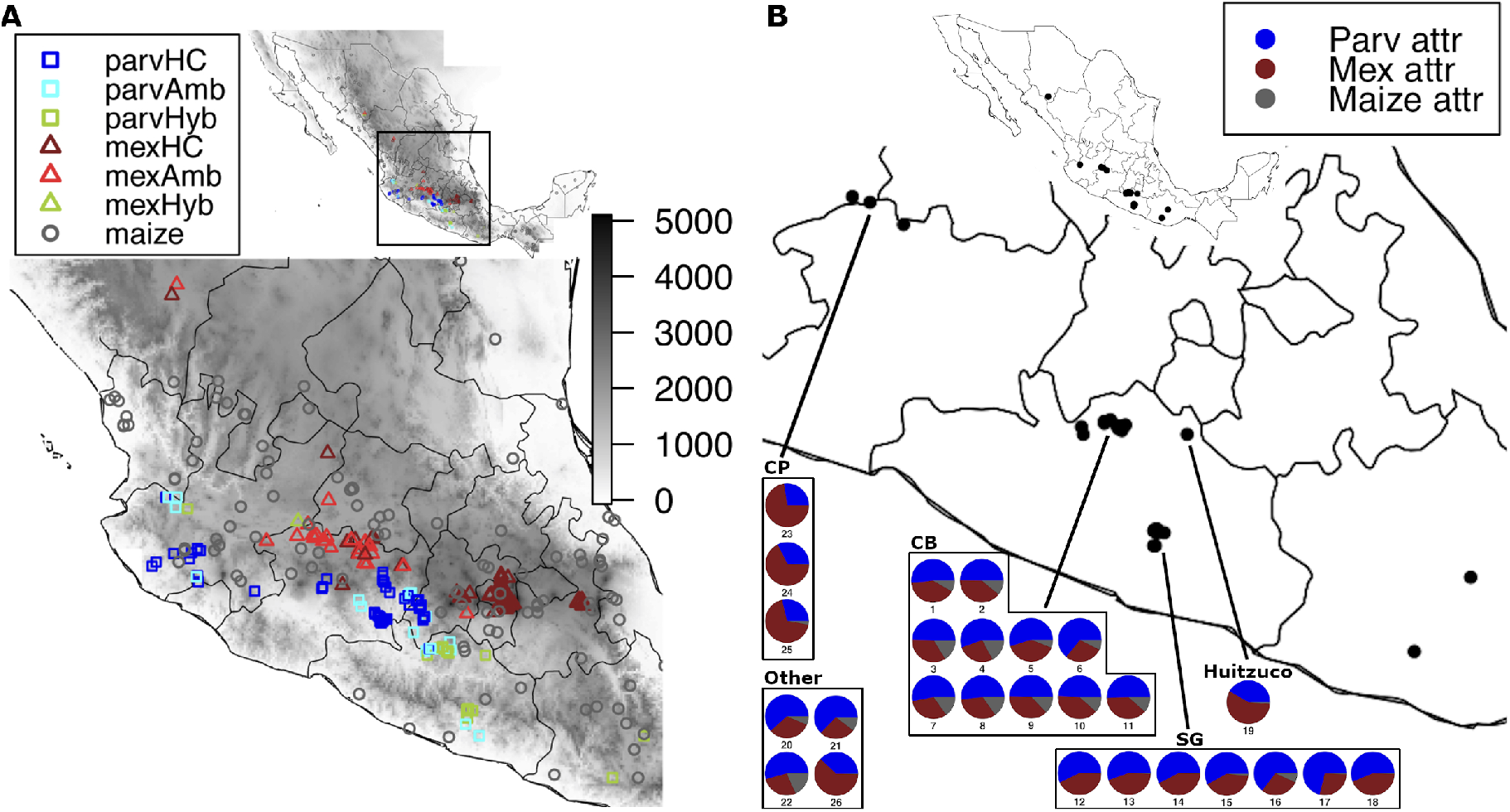
Samples plotted on Mexico. A) All 1,344 parviglumis, mexicana, and maize individuals in the published 967-SNP data set along with altitudes from the National Aeronautics and Space Administration’s (NASA’s) Shuttle Radar Topography Mission(SRTM) data set (Farr *et al*. (2007)). B) All hybrid populations in the published 967-SNP data set. Each pie chart represents a hybrid population and each color within a pie chart represents the global attribution to parviglumis, mexicana, or maize averaged over all individuals in the population based on the STRUCTURE q-matrix.

**Figure 2.**
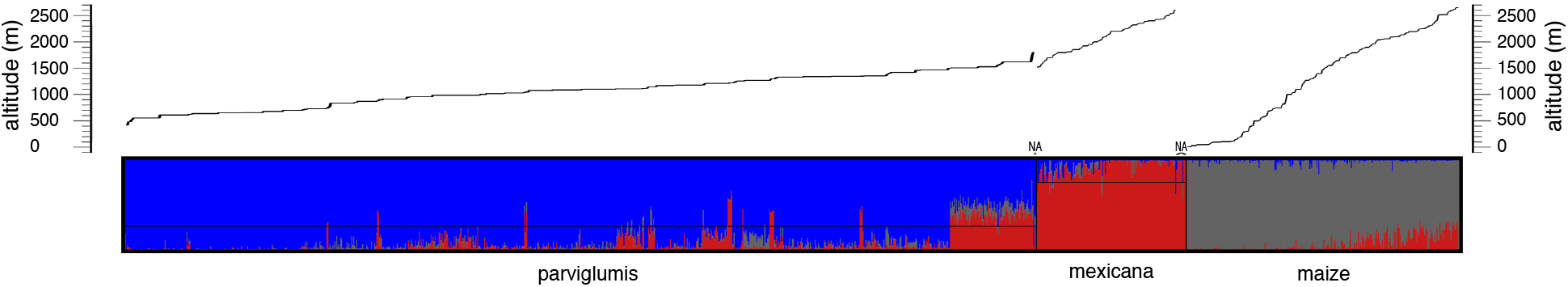
STRUCTURE q-value attributions for all 1,344 individuals in the published 967-SNP data set for k=3. Each vertical line shows the attribution of an individual to the three subspecies based on per loci group assignments for all loci in the data set. Horizontal lines were added to indicate the cutoff for identifying high-confidence hybrids. Plants within each taxonomic group are ordered from lowest to highest altitude with individuals with unknown to the right of the highest altitude individuals. Altitude is plotted above the distruct plot. This plot was generated using the program distruct (version 1.1) (Rosenberg *et al*. (2002)).

In order to determine the consistency of hybridization within populations we also performed a STRUCTURE analysis of known parviglumis, mexicana, and hybrid populations from our generated 820-SNP data set, which has deeper population sampling. Consistent with our previous analysis, we observed clear k-groups consisting of parviglumis and mexicana with hybrids observed between these taxa. In the most supported model of k=3 groups the parviglumis population Purificación was assigned to its own group, with remaining parviglumis populations assigned to a second group. All mexicana populations formed a clear third group. The Chilpancingo parviglumis population was the only clear hybrid population, showing appreciable ancestry in both the mexicana and primary parviglumis groups (Fig. S2). At higher k values, the Chilpancingo population continued to show admixture. Importantly, individuals within populations, with one exception in Ayotlan, showed consistent patterns of ancestry. (Fig. S2)

### 4.2 Confirming Hybrid Identity

It is a known issue that signals of admixture from STRUCTURE in geographically intermediate populations can be caused by Isolation By Distance (IBD) rather than hybridization (Lawson *et al*., 2018). We therefore validated patterns of population structure using the R package conStruct, which generates a STRUCTURE-like q-matrix while explicitly controlling for IBD (Bradburd *et al*., 2018). Using the spatial model in conStruct and a randomly subsampled set of parviglumis, mexicana, and hybrid individuals, the signal of hybridization remained clear and overall patterns of ancestry attribution were consistent with STRUCTURE results (Fig. 3).

**Figure 3.**
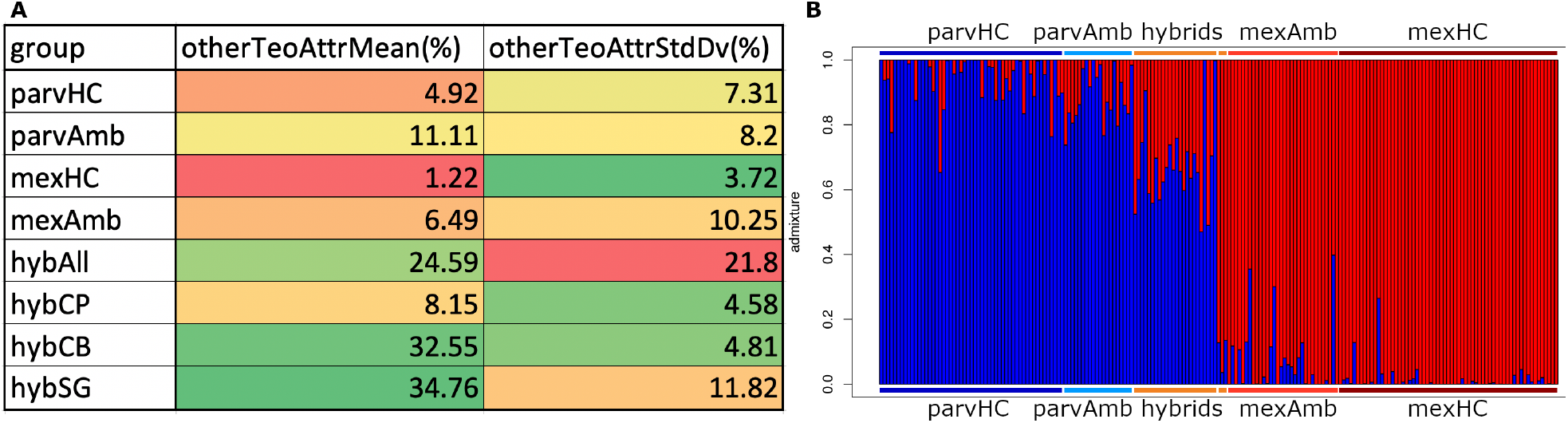
A) Percent conStruct q-value attributions to the teosinte that does not match an individual’s taxonomic description, averaged among groups with standard deviations. B) The highest likelihood STRUCTURE-like bar plot generated using the spatial model of conStruct at group size two. Each vertical line shows the attribution of an individual to parviglumis (blue) and to mexicana (red) based on per loci group assignments for all loci in the data set. Individuals within bar plots are ordered by their group as follows: high-confidence parviglumis, ambiguous parviglumis, South Guerrero hybrids, Central Balsas hybrids, ungrouped parviglumis hybrids, Central Plateau hybrids, ungrouped mexicana hybrids, ambiguous mexicana, and high-confidence mexicana. Colored horizontal bars above and below the plot represent these groups.

To further verify hybrid identity we identified four highly differentiated SNPs between parviglumis and mexicana within a large inversion (*Inv4m*) that largely distinguishes the two subspecies (Pyhajarvi *et al*., 2013) and used them to determine the inversion haplotype of all parviglumis, mexicana, and hybrid individuals. The five possible inversion states were P for parviglumis type, M for mexicana type, H for heterozygous type, R for recombinant type, and N for missing data type. Almost all high-confidence parviglumis individuals were P, with no H individuals (Fig. 4). High-confidence mexicana had 78.9% M and 21.1% R individuals, but no H individuals. Ambiguous mexicana and parviglumis included a small number of R and H plants, but otherwise showed high fidelity to their respective taxa. High confidence hybrids showed a much higher percentage of heterozygous (H) haplotypes (13.4%). Inversion haplotypes were plotted across the elevation gradient, revealing a very steep cline between parviglumis and mexicana with heterozygous hybrids in the center of the teosinte distribution (Fig. S9).

**Figure 4.**
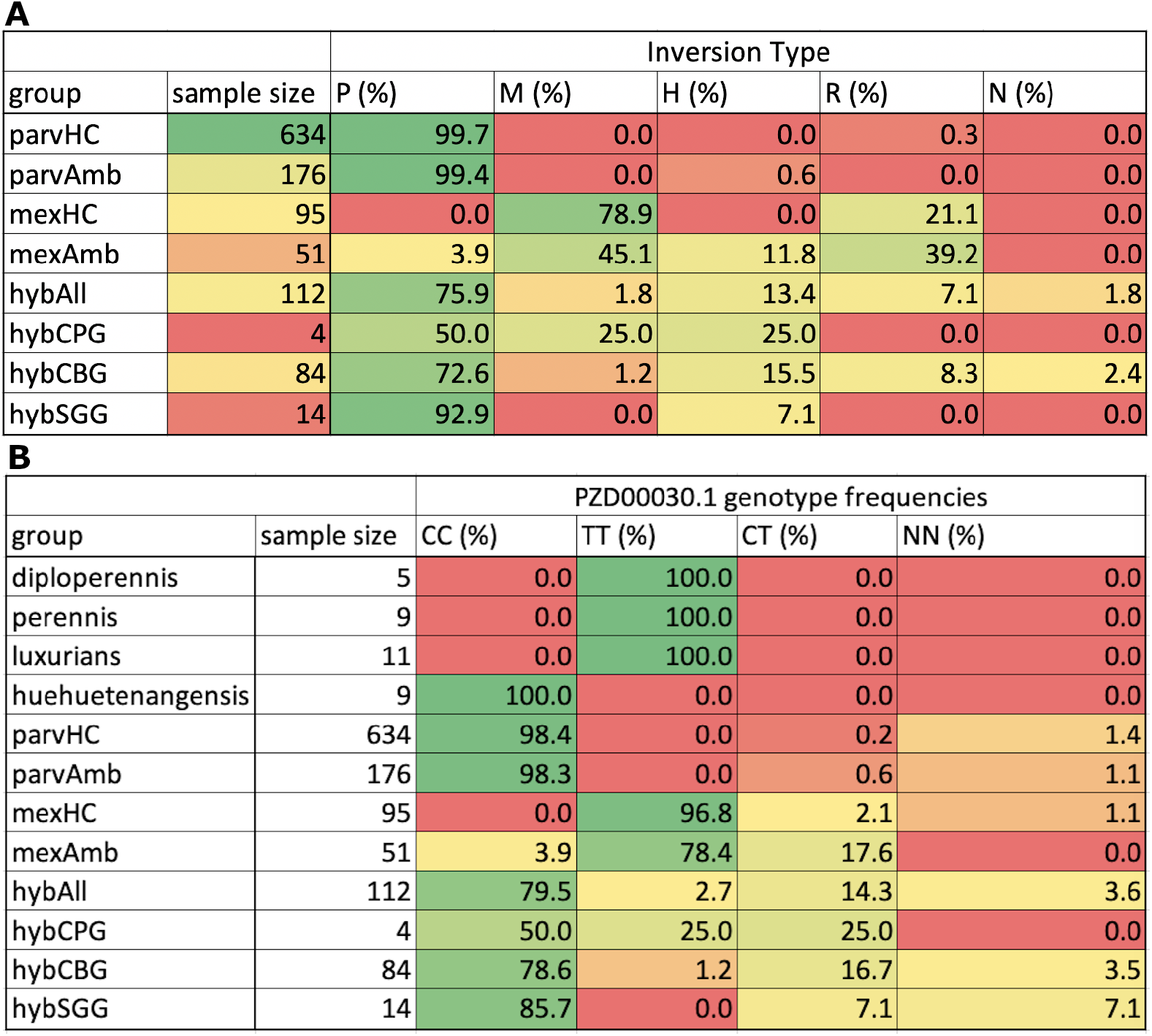
*Inv4m* haplotype and top SNP tables A) Frequency of inversion types for *Inv4m* in high-confidence and ambiguous parviglumis and mexicana individuals, all hybrids, and each hybrid group. B) Frequency of PZD00030.1 genotypes in *Zea diploperennis, Zea perennis, Zea luxurians, Zea mays huehuetenengensis*, high-confidence and ambiguous parviglumis and mexicana individuals, all hybrids, and each hybrid group.

Finally, we evaluated a third hypothesis, that putatively hybrid individuals are not the products of IBD or hybridization, but rather are ancestral to parviglumis and mexicana, based on segregation patterns of *Inv4m* in related taxa. We assessed The SNP state for the highest *F*_*ST*_ SNP on *Inv4m* in *Zea diploperennis, Zea perennis, Zea luxurians, Zea mays* ssp. *huehuetenangensis*, parviglumis, mexicana and hybrid groups (Fig. 4) and found that T was the ancestral allele for our diagnostic SNP, as it was present in 100% of more anciently diverged teosintes (*i*.*e*., *Zea diploperennis, Zea perennis*, and *Zea luxurians*). The subspecies *Zea mays* ssp. *huehuetenangensis* and parviglumis were the only taxa with the derived allele, C, and showed very little segregation. Mexicana had primarily the ancestral allele, with most exceptions being found in ambiguously assigned mexicana. Among the hybrids, the ancestral allele was segregating at a high minor allele frequency. Collectively, these segregation patterns suggest the hybrids identified here are admixed individuals rather than an ancestral population to parviglumis and mexicana.

### 4.3 Identifying Genetically Distinct Hybrid Groups

Most high-confidence hybrids were found in three distinct regions in Mexico: the Central Plateau Group (CPG), the Central Balsas Group (CBG), and the Southern Guerrero Group (SGG) (Fig. 1). These hybrid groups varied substantially in the number of assigned individuals: 4 in the CPG, 84 in the CBG, and 14 in the SGG. Note that these do not include all admixed individuals in these areas, but rather only those that passed our stringent threshold for being labeled high-confidence hybrids. The CPG has more mexicana attribution than parviglumis, and the CBG and SGG show the opposite pattern with the CBG also showing appreciable attribution to maize (Fig. 1). Differences in the prevalence of *Inv4m* haplotypes across hybrid groups were also observed (Fig. 4). The derived parviglumis type (P) became increasingly more prevalent in hybrids in southerly hybrid zones.

In order to determine whether cohesive hybrid groups could be identified based on genotype alone we ran a Principle Component Analysis (PCA) with all parviglumis, mexicana, and hybrid samples. While STRUCTURE identifies differences in global ancestry proportions, PCA more clearly captures the genetic distance of individuals and helps reveal relationships among hybrid groups and between hybrids and parental taxa. The PCA showed clear clustering of parviglumis, mexicana, and hybrid groups (Fig. 5). This validates both our threshold for identifying high confidence hybrids as well as our grouping of said hybrids. Overall, PC1 distinguished between parviglumis and mexicana individuals with hybrids intermediate, while PC2 distinguished between different hybrid groups (Fig. 5).

**Figure 5.**
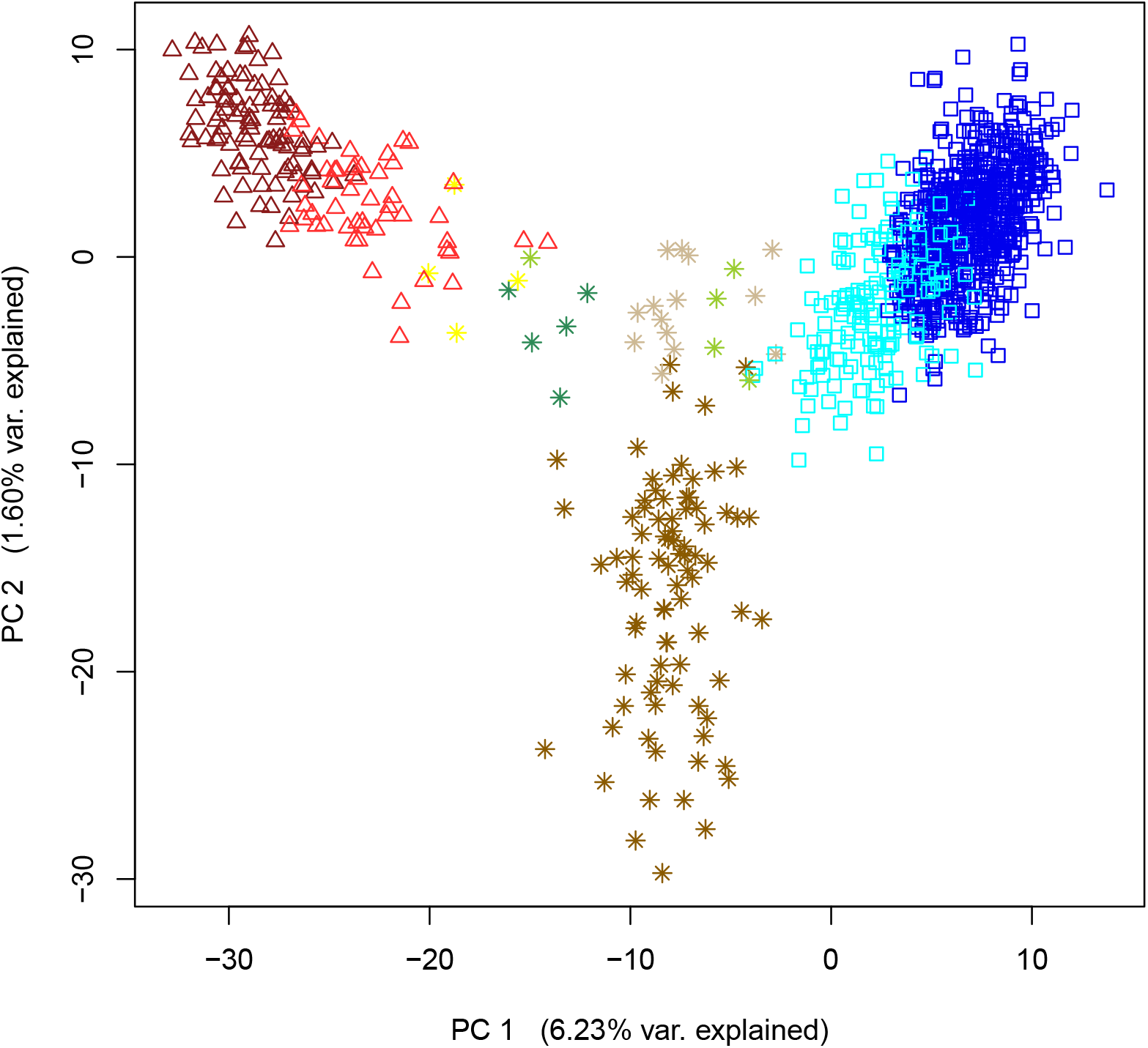
Principle Component Analysis of parviglumis, mexicana, and hybrid samples based on the R package “prcomp”. Colors and shapes correspond to taxonomic groups as follows: dark blue squares=high confidence parviglumis, light blue squares=ambiguous parviglumis, dark red triangles=high confidence mexicana, light red triangles=ambiguous mexicana, yellow asterisks=central plateau hybrid group, brown asterisks=central Balsas hybrid group, tan asterisks=South Geurrero hybrid group, seagreen asterisks=Huitzuco, and yellowgreen asterisks=other hybrids

A biplot of PC1 and PC2 revealed that among the top 10 SNPs, in terms of vector magnitude, were one SNP in a large, previously identified inversion on chromosome 3 (Romero Navarro *et al*., 2017a), five SNPs in *Inv4m*, a large inversion on chromosome four (Pyhajarvi *et al*., 2013), and one SNP located in the ZMM4 gene, a MADS-box gene which affects flowering time and development ((Danilevskaya *et al*., 2008); Figs S3 & S4). In fact, four of the five SNPs in *Inv4m* were the top four SNPs in the entire data set by this measure.

The high degree of differentiation of the CBG from parviglumis, mexicana, and other hybrid groups along PC2 could possibly be attributed to the maize ancestry observed in our STRUCTURE analysis; To assess this the PCA was broadened to include Mexican maize (Fig. 5). We found that the CBG did not overlap with maize (Fig. S5), but was closer to maize than other hybrid groups.

### 4.4 Environmental Characteristics of Hybrid Zones

In order to assess hybrid zone habitats we used the geographic coordinates of the individuals in hybrid groups to extract mean annual temperature and precipitation data with the R package, “dismo” (Hijmans *et al*., 2005) (Fig. 6). We also included all high-confidence non-admixed parviglumis and mexicana individuals for comparison. With mexicana’s habitat being cool and dry and parviglumis’ habitat being warm and wet we expected hybrids to have intermediate environments relative to parental taxa based on their intermediate altitudes. Together, these data show that the CPG’s environment is intermediate in altitude and temperature, and mexicana-like in its precipitation. The CBG’s habitat is intermediate in altitude and temperature, and parviglumis-like in its precipitation. Finally, the SGG’s environment is intermediate in temperature, and parviglumis-like in its altitude and precipitation (Fig. 6).

**Figure 6.**
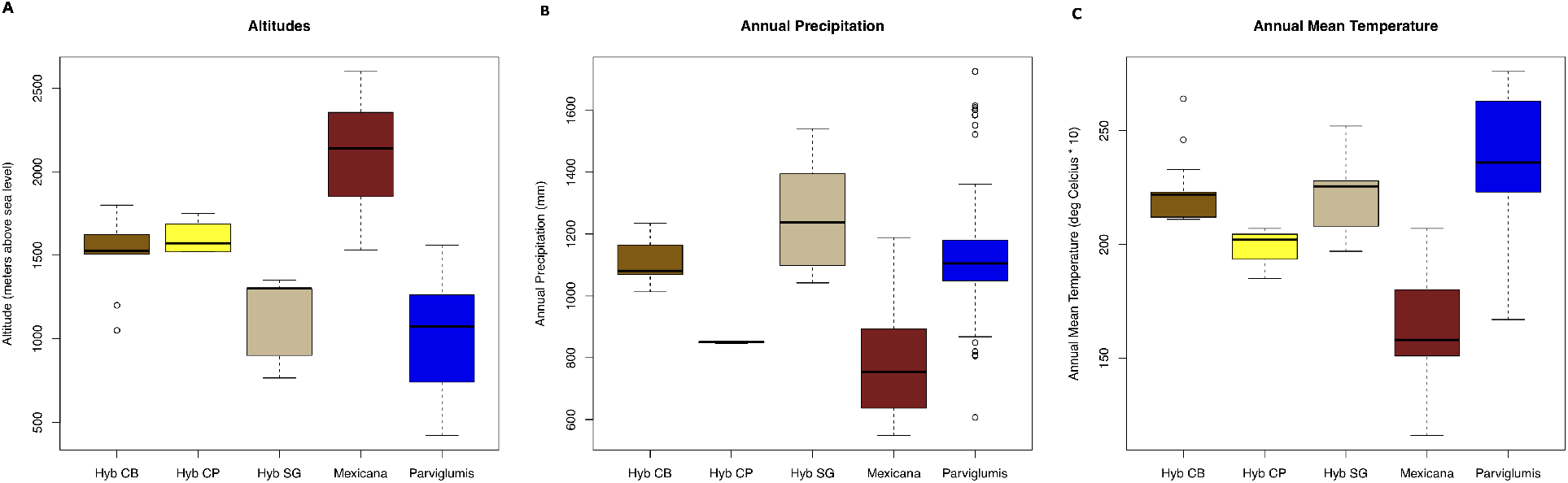
Boxplots of environmental variables for hybrid groups, parviglumis, and mexicana samples A) altitudes in meters above sea level, B) Annual precipitation in millimeters C) annual mean temperature in degrees Celsius times ten.

### 4.5 Evidence for Genomic Architectures of Hybridization

To further clarify the genetic distinctness of hybrid groups and their relationship to the major races of parviglumis (races Balsas & Jalisco) and mexicana (races Central Plateau & Chalco), we compared them using Wright’s global pairwise *F*_*ST*_ (Wright, 1952). Hybrid individuals from SGG and CBG were very similar to each other and somewhat differentiated from the CPG (Fig. S6). Furthermore, all hybrid groups were most similar to the Balsas parviglumis race and the Central Plateau mexicana race indicating a possible single origin.

To determine whether distinct patterns of hybridization could be found in different groups, we compared pairwise per-locus *F*_*ST*_ values of hybrid groups to both parviglumis and mexicana. The differentiation of the CBG and SGG groups relative to parviglumis was generally low across SNPs and the groups were similar to each other, with the CPG having many more obviously differentiated loci; this pattern was seen genome-wide with few exceptions, which matches expectations based on our estimates of global ancestry proportions. In contrast, we found substantial variation in per-locus *F*_*ST*_ of hybrid groups relative to mexicana across chromosomes, suggesting hybrid groups have different hybrid architectures of mexicana alleles and/or different sources of mexicana ancestry; this was even true comparing the CBG to the SGG, which are otherwise very similar. For example, on chromosome 1 the CBG had much more mexicana-like allele frequencies despite having similar global attributions to mexicana as the SGG. Similarly, regions of chromosome 4 showed high *F*_*ST*_ markers for the SGG relative to mexicana as compared to the CBG (Figures S7 & S8). Whether these distinct patterns of parental subspecies ancestry are adaptive in hybrids is uncertain and will require higher density data for assessing signatures of selection across hybrid zones.

## 5 Discussion

## 5.1 Hybrids May be Adapted to Intermediate Altitudes

We used a low-density genomic SNP data set to identify 112 parviglumis-mexicana hybrids, 107 in three genetically-distinct allopatric hybrid groups in Central Mexico. Hybrids were identified based on STRUCTURE’s q-matrix using a strict threshold, as were non-admixed populations of parviglumis and mexicana. Additionally, we ran a STRUCTURE analysis of deeply sampled populations using an 820-SNP data set, which showed consistency in admixture proportions across individuals at the population level.

PCA confirmed hybrid groupings and a biplot identified SNPs in large inversions on chromosomes three and four as well as a SNP within the gene ZMM4 as being among the most informative SNPs in differentiating between teosinte subspecies and between hybrid populations. Eigenvectors of these SNPs lie primarily along PC1, but are also influenced by PC2 (Fig. S3). ZMM4 and the inversion on chromosome 3 have both been associated with flowering time (Romero Navarro *et al*., 2017b; Portwood *et al*., 2019). Furthermore, the four most important SNPs for distinguishing parviglumis, mexicana, and hybrids were within the *Inv4m* inversion on chromosome 4 (Fig. S4). *Inv4m* has been linked to traits related to adaptation to high altitudes including flowering time, pigmentation, and macrohair development (Lauter *et al*., 2004; Pyhajarvi *et al*., 2013; Hufford *et al*., 2013; Romero Navarro *et al*., 2017a; Fustier *et al*., 2019). Adaptive variation therefore distinguishes parviglumis, mexicana, and hybrids along the altitudinal gradient of western Mexico, and contributes, along with the neutral effects of genetic drift, to their differentiation. These findings are consistent with previous literature which identified clear signatures of local adaptation in parviglumis and mexicana (Wilkes, 1967; Doebley, 1984; Pyhajarvi *et al*., 2013; Fustier *et al*., 2019, 2019).

### 5.2 Hybrids May Share a Common Origin

*F*_*ST*_ comparisons revealed that one race of each parent subspecies shows greatest similarity to all hybrids, suggesting a possible single origin for all three hybrid groups. It is also possible that these hybrid groups originated from different parental populations within the same races. Global *F*_*ST*_ results also showed greater similarity between the SGG and the CBG than either have with the CPG. This could suggest more recent divergence between the SGG and the CBS relative to the CPG, post-hybridization introgression with nearby teosinte, local adaptation to more similar environments, or some combination of these potential phenomena.

### 5.3 These Populations are Composed of True Hybrids

Although many groups have worked with these hybrid populations (e.g., (Aguirre-Liguori *et al*., 2017; Fustier *et al*., 2017)), none have suggested potential hybrid zone dynamics between mexicana and parviglumis even when admixture between the subspecies was detected (Fukunaga *et al*., 2005; Pyhajarvi *et al*., 2013; Aguirre-Liguori *et al*., 2019; Fustier *et al*., 2019; van Heerwaarden *et al*., 2011). Perhaps no conclusion has previously been made due to the alternative explanations for genetic intermediates we have shown here to have little support. The density and broad geographic distribution of the samples analyzed here helped clarify that there are distinct regions in which hybridization occurs.

In order to make a strong case that these plants are true hybrids, alternative hypotheses needed to be ruled out. For example, putative hybrids in intermediate geographic locations could be the products of IBD when their identification relies on STRUCTURE (Lawson *et al*., 2018). We implemented the R package conStruct to generate a STRUCTURE-like q-matrix while explicitly controlling for IBD and demonstrated that the signal of admixture did not disappear (Bradburd *et al*., 2018). Putative hybrids could also represent an ancestral taxon from which parviglumis and mexicana were derived. We analyzed the distribution of inversion types and allele frequencies of *Inv4m* across altitudinal clines and taxonomic groups, showing a pattern consistent with hybridization rather than an ancestral relationship relative to parviglumis and mexicana. One implication of this is that it resolves the phylogeny of parviglumis such that it is now monophyletic if one excludes maize (Fukunaga *et al*., 2005), which was domesticated from parviglumis (Matsuoka *et al*., 2002). A second implication is that future teosinte studies can avoid hybrid populations when the goal is to characterize parviglumis and mexicana and utilize these populations when the goal is to study hybridization dynamics and hybrid zones.

### 5.4 Hybrids Have Unique Environments and Potentially Unique Population Dynamics

While all of our putative hybrid zone environments are generally intermediate between those of the subspecies, there are marked distinctions in terms of temperature, precipitation, and altitude. A general pattern was observed in which the level of shared genetic ancestry between hybrids and both parviglumis and mexicana was correlated with the extent to which their habitats were parviglumis-like or mexicana-like in nature. Although we measured habitats in altitude, temperature, and precipitation, many other environmental characteristics may vary across these regions that were not measured directly here and could potentially be driving differences in hybrid genotypes and phenotypes via local adaptation (Korner, 2007).

Hybrid populations may also be characterized by distinct population dynamics (*e*.*g*., falling within tension zones, neutral intergadation zones, or selection-dominated zones (Barton and Hewitt, 1985)). The CPG is the only group that appears, based on our sampling, to reside in a current range overlap between parviglumis and mexicana. Therefore, while it could be a selection-dominated zone, it is the only group that has a reasonable probability of representing a tension zone or a neutral intergradation zone. As for the CBG and SGG, nearby parviglumis populations are documented, but nearby mexicana populations are not known. Ecological niche modeling suggests the nearest mexicana samples may be over 100km away from the CBG and even further from the SGG (Hufford *et al*., 2012). Although unlikely, it is possible that rare long-range dispersal could be contributing to ongoing hybridization in the CBG (Cain *et al*., 2000; Nathan, 2006). More likely scenarios could be that the CBG and SGG were formed at a time when mexicana was more broadly distributed, that nearby and unsampled mexicana individuals are contributing to admixture, or that hybrid groups originated in one or more hybrid zones and subsequently spread across Mexico. Regardless of how they arrived in their current location, CBG and SGG may lack ongoing gene flow from mexicana. We therefore speculate that both are selection-dependent zones, and, given their physical isolation could be on the path to speciation (Barton and Hewitt, 1985; Abbott *et al*., 2013; Abbott and Brennan, 2014; Abbott *et al*., 2008; Rieseberg *et al*., 1998; Heiser *et al*., 1969).

### 5.5 Teosinte Hybrids are a Promising Model Study System

Hybrid zone dynamics have been studied in a diverse set of species with each system having its own particular strengths for biological inference. For example, sunflowers (*Helianthus* spp.) represent one of the most established model systems for the study of hybridization and have generated substantial insight into the processes of hybrid speciation and hybrid zone dynamics. Three distinct species, each with a unique ecological niche, have resulted from repeated hybridization of the same parental sunflower species (*Helianthus annuus* x *H. petiolaris*) Gross and Rieseberg (2004). Additionally, in natural hybrid zones of these parental species, analysis of the genetic architecture of hybridization has shown clear barriers to gene flow between the taxa in certain chromosomal regions, likely due to genetic isolation factors Rieseberg *et al*. (1999). *Senecio squalidus* is another established model system where an altitude-based hybrid zone on Mount Etna, Sicily has resulted in hybrid speciation, with the novel species subsequently becoming invasive throughout England following escape from cultivation in the Oxford Botanic Garden (Abbott *et al*., 2008). The *Senecio* system is therefore excellent for studying how hybrid speciation can lead to novel and broad adaptation and potential invasiveness.

The teosinte hybrid groups identified in this study are genetically distinct and grow in varying environments. Altitude is the primary distinguishing feature of the ranges of their ancestral subspecies, parviglumis and mexicana, which have distinct morphological features that suggest local adaptation (Wilkes, 1967; Doebley, 1984; Hufford *et al*., 2012; Fukunaga *et al*., 2005; Aguirre-Liguori *et al*., 2017; Fustier *et al*., 2017; Aguirre-Liguori *et al*., 2019). We found evidence of distinct architectures of hybridization in hybrid groups. Differential retention of parental haplotypes across hybrid groups may suggest local adaptation in each distinct region, a possibility that should be further explored when signatures of selection can be assessed in a data set with higher marker density. Teosinte, as a system, therefore has excellent potential for exploring adaptive architectures of hybridization across multiple, independent hybrid zones that are distributed across a broad latitudinal gradient.

In conclusion, we have identified and confirmed a set of 112 hybrids, mostly clustered in three allopatric, genetically and environmentally distinct hybrid groups (the CPG, CBG and SGG), which were confirmed by PCA clustering. We provided evidence that these individuals are true hybrids and not ancestral to parviglumis and mexicana, or products of IBD using PCA, additional structure runs with an 820-SNP data set, inversion analyses using *Inv4m*, and a conStruct analysis. In the future, we would like to use a denser genomic data set to look for genomic signals of local adaptation and genomic architectures of hybridization. With the potential for local adaptation, variable hybrid zone dynamics, and differential architectures of hybridization in all three populations, we present this parviglumis-mexicana hybrid system as a great, natural model system for studying hybridization and hybrid zones.

## 6 Acknowledgements

We would like to thank Jeffrey Ross-Ibarra of the University of California Davis for his advice on this project. SNP generation was paid for by NSF grant IOS 1238014 to JFD. While at Iowa State University stipend support for D.E.H. was provided by the Plant Sciences Institute and start-up funds to MBH from Iowa State University. D.E.H. was funded, during the editing and revision process, by the USDA Agricultural Research Service Research Participation Program of the Oak Ridge Institute for Science and Education (ORISE) through an interagency agreement between the U.S. Department of Energy (DOE) and USDA Agricultural Research Service (contract number DE-AC05-06OR23100). Mention of trade names or commercial products in this article is solely for the purpose of providing specific information and does not imply recommendation or endorsement by the USDA, DOE, or ORISE. USDA is an equal opportunity provider and employer. No funding bodies had any role in study design, data collection, analysis and interpretation, writing, or the decision to submit the work for publication.

## 7 Author Contributions

DEH and MBH designed research goals and methods. JFD and JJSG generated SNP data. DEH and KK analyzed data. DEH, KK, and MBH interpreted results. DEH wrote the paper.

