## Supplemental Materials for "Evidence for Multiple Teosinte Hybrid Zones in Central Mexico"

February 11, 2021

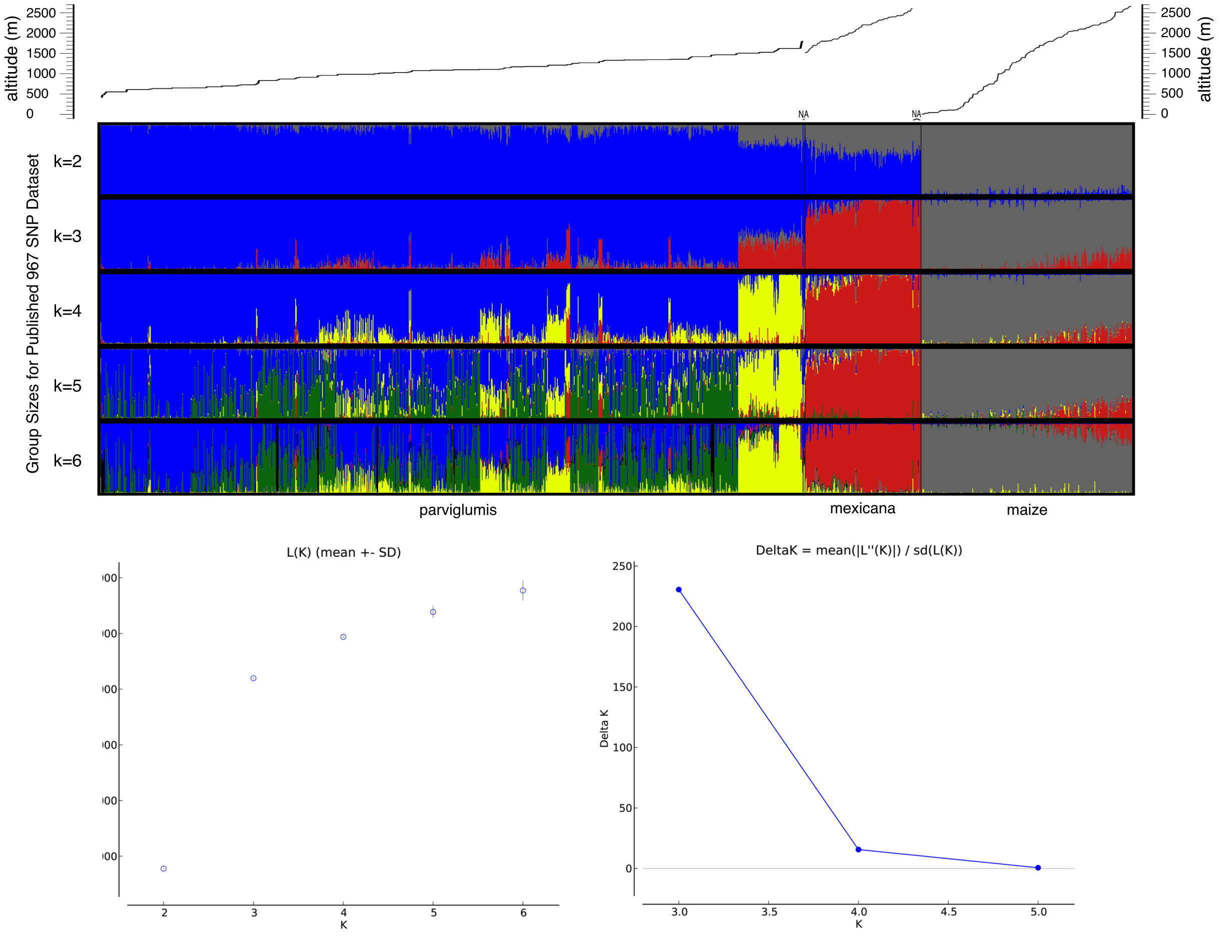


Figure S1: STRUCTURE q-value attributions for all 1,344 individuals in the published 967 SNP data set for all k values from 2 to 6. Each vertical line shows the attribution of an individual to the three subspecies based on per loci group assignments for all loci in the data set. Plants within each taxonomic group are ordered from lowest to highest altitude with individuals with unknown to the right of the highest altitude individuals. Altitude is plotted above distruct plots.

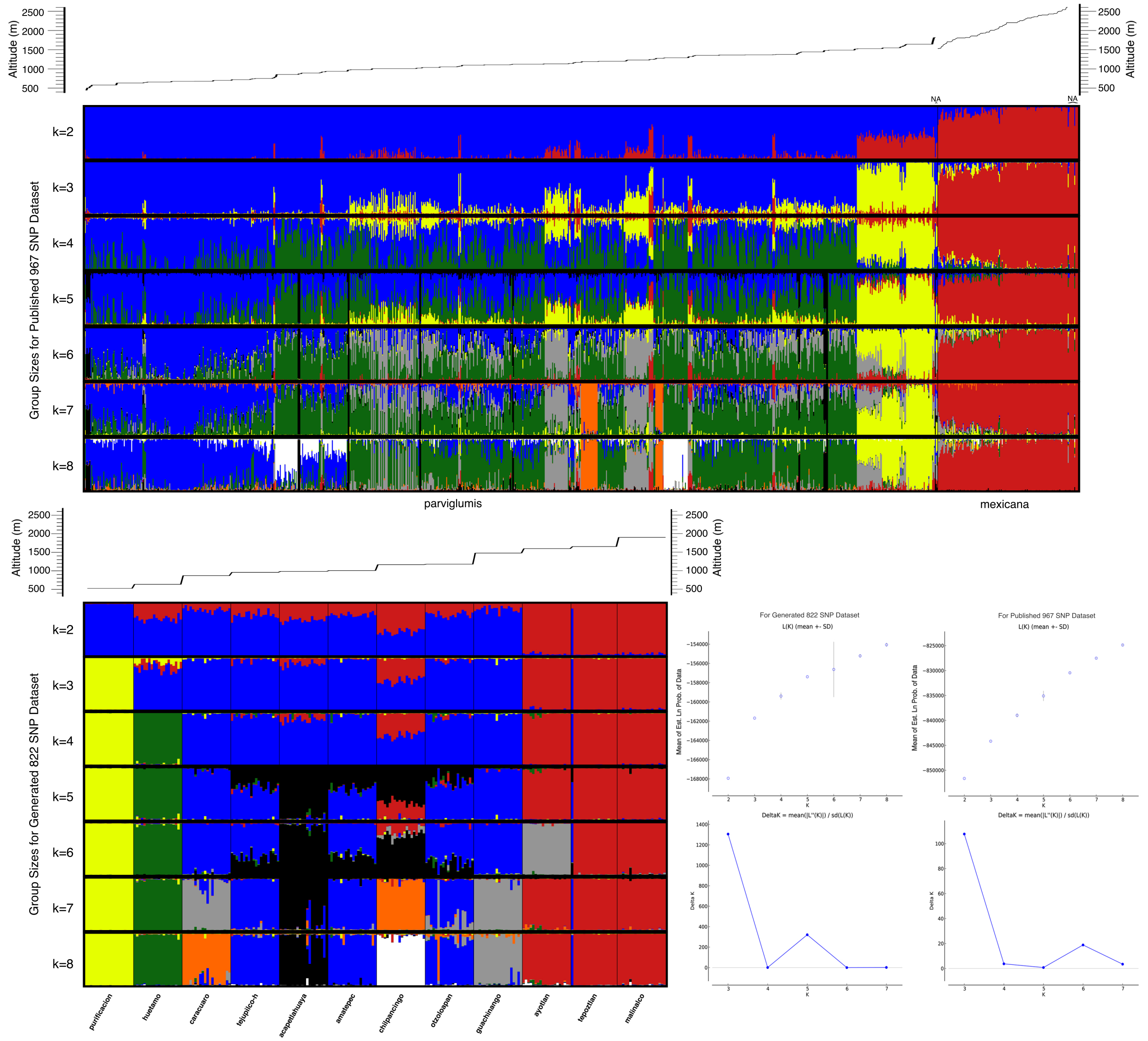


Figure S2: STRUCTURE q-value attributions for all 1,344 individuals in the published 967 SNP data set and all 232 individuals in the generated 822 SNP data set for all k values from 2 to 8. Each vertical line shows the attribution of an individual to the three subspecies based on per loci group assignments for all loci in the data set. Plants within each taxonomic group are ordered from lowest to highest altitude with individuals with unknown to the right of the highest altitude individuals. Altitude is plotted above distruct plots.

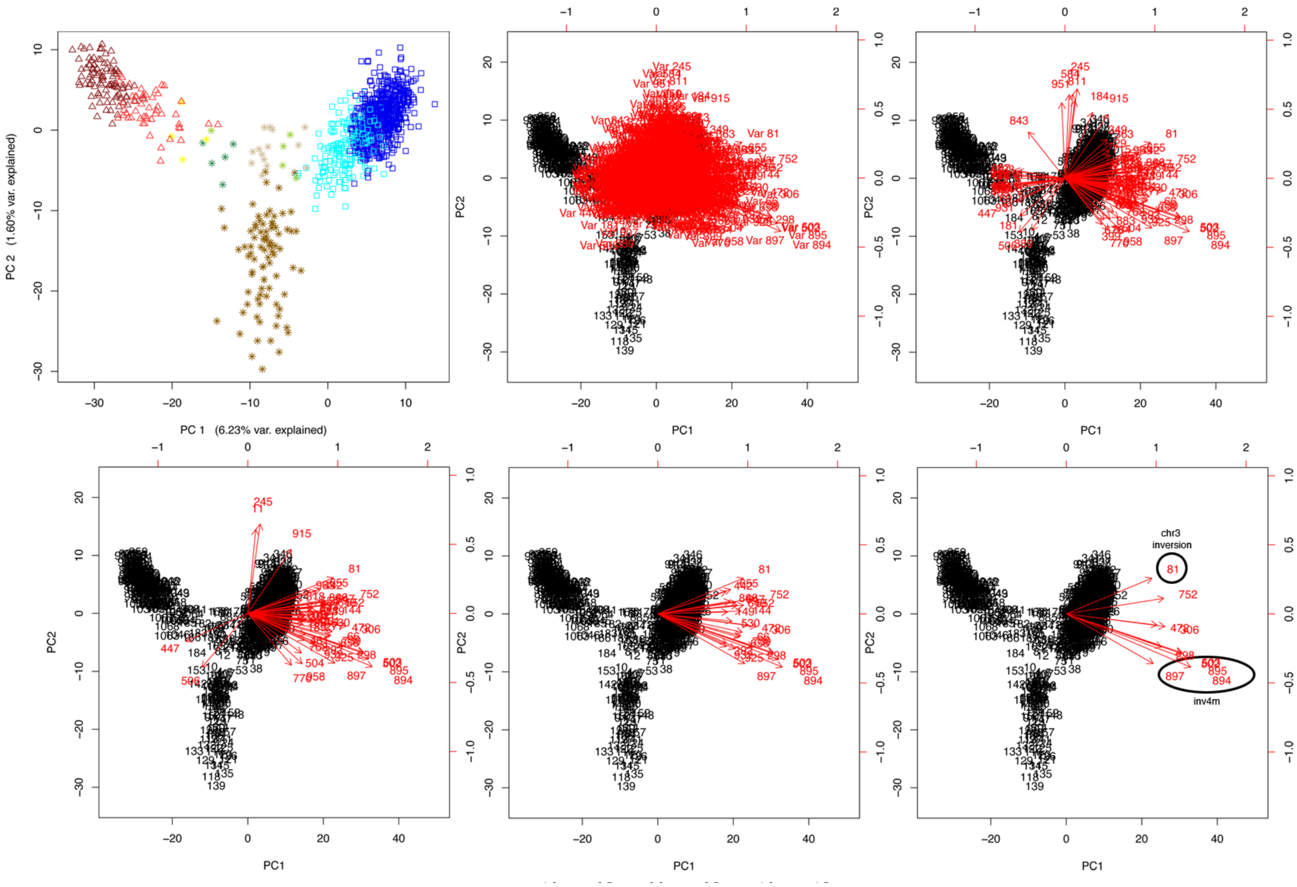


Figure S3: From left to right and top to bottom: 1) the Principle Component Analysis of teosinte samples. Colors and shapes correspond to taxonomic groups as follows: dark blue squares=high confidence parviglumis, light blue squares=ambiguous parviglumis, dark red triangles= high confidence mexicana, light red triangles= ambiguous mexicana, yellow pluses=central plateau hybrid group, brown pluses =central Balsas hybrid group, tan pluses=South Geurrero hybrid group, seagreen pluses=Huitzuco, and yellowgreen pluses =other hybrids. 2) Biplot with all SNPs 3) Biplot with top 100 SNPs in terms of vector magnitude on PC1 and PC2 4) Biplot with top 50 SNPs 5) Biplot with top 25 SNPs 6) Biplot with top 10 SNPs. Biplots were made using the R package ”biplot”.

| markerName | rank | PCAvecMag | markerNum | PCAxCoord | PCAyCoord | PCAangle | PCAprimaryCont | notes |
|---|---|---|---|---|---|---|---|---|
| PZB01116.2 | 100 | 0.66008396 | 843 | -0.5099828 | 0.4190804 | -39.411797 | parv_mex_hyb | |
| PZA00565.3 | 99 | 0.66065551 | 389 | -0.4615568 | -0.4726849 | 45.6824482 | Ambiguous | |
| PZA03312.2 | 98 | 0.66121498 | 676 | 0.54454546 | -0.3750673 | -34.557962 | parv_mex_hyb | |
| PZB01114.3 | 97 | 0.66438307 | 839 | 0.6449365 | -0.1595674 | -13.896803 | parv_mex_hyb | |
| PZA03774.9 | 96 | 0.66480997 | 715 | 0.63071528 | 0.21016787 | 18.4291827 | parv_mex_hyb | |
| PZA00588.2 | 95 | 0.66482517 | 399 | 0.51527115 | -0.4201049 | -39.190658 | parv_mex_hyb | |
| PZA00497.1 | 94 | 0.66584205 | 338 | -0.6628379 | 0.06317885 | -5.4447387 | parv_mex_hyb | |
| PZA00006.14 | 93 | 0.66601055 | 29 | 0.61570041 | 0.25393514 | 22.4128679 | parv_mex_hyb | |
| PZA03047.22 | 92 | 0.66678882 | 613 | 0.65967493 | -0.0971407 | -8.3769139 | parv_mex_hyb | |
| PZA00562.4 | 91 | 0.66837745 | 388 | 0.66831804 | -0.0089116 | -0.7639581 | parv_mex_hyb | |
| PZA00337.3 | 90 | 0.67117392 | 224 | 0.67116541 | -0.0033794 | -0.2884898 | parv_mex_hyb | |
| PZA00096.26 | 89 | 0.67451312 | 69 | -0.6590306 | -0.1436894 | 12.2997978 | parv_mex_hyb | |
| PZA00017.1 | 88 | 0.67695557 | 36 | 0.6729755 | -0.0732996 | -6.2160751 | parv_mex_hyb | |
| PZA03786.2 | 87 | 0.67834555 | 736 | 0.65287683 | -0.1841318 | -15.750125 | parv_mex_hyb | |
| PZA00547.18 | 86 | 0.67900671 | 384 | -0.6741662 | -0.0809324 | 6.84549665 | parv_mex_hyb | |
| PZA00504.2 | 85 | 0.68520094 | 349 | 0.5872405 | 0.35305654 | 31.01483 | parv_mex_hyb | |
| su1.7 | 84 | 0.6854072 | 951 | -0.0446502 | 0.68395131 | -86.264892 | parv_mex_hyb | |
| PZD00020.2 | 83 | 0.68977031 | 881 | 0.67723756 | 0.13089066 | 10.9387668 | parv_mex_hyb | |
| PZA00593.2 | 82 | 0.69370494 | 404 | -0.6910281 | -0.0608827 | 5.0350196 | parv_mex_hyb | |
| PZA03311.4 | 81 | 0.69440789 | 674 | 0.66756079 | -0.1912195 | -15.984103 | parv_mex_hyb | |
| Fea2.1 | 80 | 0.69800624 | 12 | -0.6980013 | -0.002626 | 0.21555625 | parv_mex_hyb | |
| PZD00076.1 | 79 | 0.70175345 | 931 | -0.6690901 | -0.2116043 | 17.5499139 | parv_mex_hyb | |
| PZA00449.2 | 78 | 0.70315684 | 299 | 0.69809752 | -0.0841986 | -6.8773091 | parv_mex_hyb | |
| PZA03305.6 | 77 | 0.70823403 | 670 | -0.7079396 | 0.02042071 | -1.6522544 | parv_mex_hyb | |
| PZA00273.1 | 76 | 0.70945272 | 184 | 0.38537508 | 0.59565863 | 57.0982294 | CB_others | genic sequence, function unknown (mRNA identified) |
| PZB01463.4 | 75 | 0.71053366 | 863 | 0.69408477 | -0.1520014 | -12.352506 | parv_mex_hyb | |
| PZD00076.4 | 74 | 0.71194457 | 933 | -0.7083423 | -0.0715277 | 5.76612288 | parv_mex_hyb | |
| PZA00263.14 | 73 | 0.71290404 | 181 | -0.6261518 | -0.3408316 | 28.5606674 | parv_mex_hyb | |
| PZD00077.7 | 72 | 0.71375691 | 936 | -0.7114914 | -0.056824 | 4.56629366 | parv_mex_hyb | |
| PZA03223.3 | 71 | 0.71559447 | 657 | -0.7118773 | 0.07284347 | -5.8425072 | parv_mex_hyb | |
| PZB00498.2 | 70 | 0.71704129 | 811 | 0.13789753 | 0.70365651 | 78.9121171 | CB_others | sbp1, SBP-domain protein1 |
| PZA02958.17 | 69 | 0.72165338 | 566 | 0.71260061 | -0.1139472 | -9.0848807 | parv_mex_hyb | |
| PZA00261.6 | 68 | 0.72384942 | 180 | -0.7235806 | 0.01972667 | -1.5616443 | parv_mex_hyb | |
| PZD00067.2 | 67 | 0.7248871 | 918 | -0.7240235 | 0.03537307 | -2.7970332 | parv_mex_hyb | |
| PZA00692.5 | 66 | 0.72717817 | 460 | -0.7255991 | -0.0478954 | 3.77650989 | parv_mex_hyb | |
| PZA00307.17 | 65 | 0.72734913 | 207 | 0.72734122 | -0.0033923 | -0.2672248 | parv_mex_hyb | |
| PZD00067.3 | 64 | 0.72862747 | 919 | -0.7277852 | 0.03502345 | -2.7551386 | parv_mex_hyb | |
| PZA00710.16 | 63 | 0.73065075 | 469 | -0.7233128 | -0.1032914 | 8.12707576 | parv_mex_hyb | |
| PZA00424.1 | 62 | 0.73093056 | 284 | 0.63087568 | -0.3691278 | -30.332075 | parv_mex_hyb | |
| PZA00425.4 | 61 | 0.73210721 | 285 | 0.72913967 | 0.06585063 | 5.16054193 | parv_mex_hyb | |
| PZB00416.2 | 60 | 0.73223974 | 805 | 0.70694644 | -0.1907925 | -15.103301 | parv_mex_hyb | |
| PZA00525.2 | 59 | 0.73257644 | 363 | 0.65847652 | 0.32105595 | 25.9926571 | parv_mex_hyb | |
| PZB01110.2 | 58 | 0.73601807 | 828 | 0.72831492 | 0.10620725 | 8.29673491 | parv_mex_hyb | |
| PZB00092.1 | 57 | 0.73644381 | 761 | 0.71522542 | -0.1755052 | -13.787084 | parv_mex_hyb | |
| PZD00020.4 | 56 | 0.7439287 | 883 | 0.67791831 | -0.3063604 | -24.318852 | parv_mex_hyb | |
| te1.4 | 55 | 0.74867027 | 957 | 0.73098596 | 0.16176125 | 12.4780027 | parv_mex_hyb | |
| PZA03001.18 | 54 | 0.75849863 | 584 | 0.05900849 | 0.75619982 | 85.5380967 | CB_others | probable methyltransferase PMT1 |
| PZA02844.1 | 53 | 0.75994136 | 527 | 0.75661422 | -0.0710337 | -5.3634164 | parv_mex_hyb | |
| PZA02872.1 | 52 | 0.76208797 | 537 | 0.75812589 | 0.07760934 | 5.8450079 | parv_mex_hyb | |
| PZA03335.2 | 51 | 0.76221872 | 687 | 0.74815088 | 0.14576566 | 11.0250795 | parv_mex_hyb | |
| PZB00607.2 | 50 | 0.76250606 | 818 | 0.75244677 | 0.12344777 | 9.31704847 | parv_mex_hyb | |
| PZB00145.2 | 49 | 0.7674088 | 770 | 0.60691233 | -0.4696527 | -37.734125 | parv_mex_hyb | |
| bt2.8 | 48 | 0.76803525 | 11 | 0.10939017 | 0.76020519 | 81.8116098 | CB_others | ADP-glucose pyrophosphorylase endosperm small subunit (bt2) gene |
| PZA00472.2 | 47 | 0.77118666 | 315 | 0.77111405 | -0.0105818 | -0.7862044 | parv_mex_hyb | |
| PZA00193.2 | 46 | 0.7717198 | 132 | 0.76891814 | -0.0656989 | -4.8836813 | parv_mex_hyb | |
| PZA00287.1 | 45 | 0.77456848 | 189 | 0.768111 | -0.0998089 | -7.4035758 | parv_mex_hyb | |
| PZA00514.1 | 44 | 0.787439 | 354 | 0.78696121 | -0.0274267 | -1.9960307 | parv_mex_hyb | |
| PZA01782.2 | 43 | 0.80179455 | 506 | -0.6365713 | -0.4874951 | 37.4453954 | parv_mex_hyb | |
| PZA00140.10 | 42 | 0.80681498 | 101 | 0.80668296 | -0.0145949 | -1.0365135 | parv_mex_hyb | |
| PZA00614.12 | 41 | 0.81586349 | 413 | 0.81366804 | -0.0598126 | -4.2042404 | parv_mex_hyb | |
| PZA00693.3 | 40 | 0.82052304 | 461 | 0.797036 | -0.1949145 | -13.741934 | parv_mex_hyb | |
| PZB00093.3 | 39 | 0.82249889 | 763 | 0.78549071 | -0.2439442 | -17.252853 | parv_mex_hyb | |
| PZA01725.1 | 38 | 0.82682196 | 504 | 0.7447178 | -0.3592074 | -25.749889 | parv_mex_hyb | |
| PZA00370.5 | 37 | 0.827508 | 245 | 0.17094652 | 0.80965843 | 78.0780109 | CB_others | Serine/threonine-protein kinase PEPKR2, |
| PZD00062.2 | 36 | 0.83516285 | 915 | 0.60148297 | 0.57940939 | 43.9291418 | parv_mex_hyb | |
| PZA02822.2 | 35 | 0.86911345 | 521 | 0.86890597 | -0.0189893 | -1.2519595 | parv_mex_hyb | |
| zagl1.1 | 34 | 0.88273896 | 958 | 0.75549557 | -0.4565682 | -31.145807 | parv_mex_hyb | |
| zen1.4 | 33 | 0.88562352 | 963 | 0.86180274 | 0.20402218 | 13.3189308 | parv_mex_hyb | |
| PZA00662.3 | 32 | 0.90015911 | 447 | -0.8637762 | -0.2533321 | 16.3455924 | parv_mex_hyb | |
| PZA00201.2 | 31 | 0.90154917 | 137 | 0.89977172 | 0.05658398 | 3.59842392 | parv_mex_hyb | |
| tb1.18 | 30 | 0.90166065 | 953 | 0.90149635 | 0.01721218 | 1.09381002 | parv_mex_hyb | |
| PZB01112.3 | 29 | 0.90657974 | 833 | 0.90521134 | 0.04979206 | 3.1484395 | parv_mex_hyb | |
| tb1.19 | 28 | 0.9136931 | 954 | 0.91192489 | -0.0568162 | -3.5651223 | parv_mex_hyb | |
| PZA02941.6 | 27 | 0.91392083 | 555 | 0.90945801 | 0.09020765 | 5.66454705 | parv_mex_hyb | |
| PZA00419.1 | 26 | 0.96966287 | 277 | 0.96521168 | -0.0928035 | -5.4920146 | parv_mex_hyb | |
| PZA00220.12 | 25 | 0.9715824 | 149 | 0.97143391 | 0.01698597 | 1.00174095 | parv_mex_hyb | |
| PZA00656.16 | 24 | 0.97292747 | 442 | 0.95133018 | 0.20385962 | 12.0949404 | parv_mex_hyb | |
| PZA00442.3 | 23 | 0.9881942 | 292 | 0.95235629 | -0.2637144 | -15.477801 | parv_mex_hyb | |
| PZD00076.2 | 22 | 0.99825877 | 932 | 0.95305954 | -0.2969816 | -17.307489 | parv_mex_hyb | |
| PZD00007.1 | 21 | 1.01320062 | 866 | 1.00665253 | 0.11500512 | 6.51750596 | parv_mex_hyb | |
| PZA02853.10 | 20 | 1.03358842 | 530 | 1.03145439 | -0.0663842 | -3.6824676 | parv_mex_hyb | |
| PZA03137.1 | 19 | 1.0371489 | 655 | 1.01306194 | 0.22222368 | 12.3723533 | parv_mex_hyb | |
| PZA03095.1 | 18 | 1.09119179 | 647 | 1.08604065 | 0.10590201 | 5.56941968 | parv_mex_hyb | |
| PZA03095.3 | 17 | 1.10621622 | 649 | 1.10333817 | 0.07974457 | 4.13390641 | parv_mex_hyb | |
| PZD00073.1 | 16 | 1.11748675 | 925 | 1.07147235 | -0.3173699 | -16.499279 | parv_mex_hyb | |
| PZA03258.2 | 15 | 1.155312 | 658 | 1.13574432 | -0.2117325 | -10.560209 | parv_mex_hyb | |
| PZA00214.1 | 14 | 1.15810052 | 144 | 1.15779492 | 0.02660308 | 1.31627501 | parv_mex_hyb | |
| PZA03789.4 | 13 | 1.17068934 | 739 | 1.15369207 | -0.1987666 | -9.7753732 | parv_mex_hyb | |
| PZA00092.1 | 12 | 1.18544894 | 66 | 1.17380325 | -0.1657562 | -8.0377615 | parv_mex_hyb | |
| PZA02939.6 | 11 | 1.19490194 | 552 | 1.19232308 | 0.07846227 | 3.76499062 | parv_mex_hyb | |
| PZA00109.3 | 10 | 1.23358863 | 81 | 1.1903655 | 0.32368331 | 15.2120217 | parv_mex_hyb | inversion 3, LOC103650437 uncharacterized LOC103650437 |
| PZA00717.14 | 9 | 1.26646941 | 472 | 1.26234851 | -0.1020834 | -4.6233244 | parv_mex_hyb | c-terminal zinc finger |
| PZD00030.6 | 8 | 1.2903773 | 897 | 1.2091312 | -0.4506388 | -20.440244 | parv_mex_hyb | inv4m, , zmm19, MADS-box transcription factor |
| PZA00445.18 | 7 | 1.35661318 | 298 | 1.32302827 | -0.2999925 | -12.775615 | parv_mex_hyb | MATE family efflux transporter, protects cells against oxidative stress and bile salts |
| PZB00067.2 | 6 | 1.36196548 | 752 | 1.3546643 | 0.14083532 | 5.93533443 | parv_mex_hyb | m4 protein, plays a role in both floral induction and inflorescence development |
| PZA00462.2 | 5 | 1.37556915 | 306 | 1.37078916 | -0.1145757 | -4.7778894 | parv_mex_hyb | Zea mays Vacuolar sorting protein 39 (umc2730), unknown function |
| PZA01637.3 | 4 | 1.64391291 | 502 | 1.60385907 | -0.360674 | -12.673779 | parv_mex_hyb | inv4m, pco139316 hypothetical protein |
| PZA01637.4 | 3 | 1.64796169 | 503 | 1.60995254 | -0.3518957 | -12.329529 | parv_mex_hyb | inv4m, zmm19, MADS-box transcription factor |
| PZD00030.4 | 2 | 1.73316727 | 895 | 1.68334077 | -0.4125926 | -13.771892 | parv_mex_hyb | inv4m, zmm19, MADS-box transcription factor |
| PZD00030.1 | 1 | 1.79632214 | 894 | 1.73022253 | -0.4828077 | -15.591426 | parv_mex_hyb | inv4m, zmm19, MADS-box transcription factor |

Figure S4: A table of information relating to PCA biplot results ordered by vector magnitude (from large to small) on the biplot. A green background indicates the top 10 SNPs, A light greed background indicates the top 25 SNPs and yellow indicates SNPs that primarily influence PC2, whereas the rest primarily influence PC1. Gene function information was investigated using the database MaizeGDB.

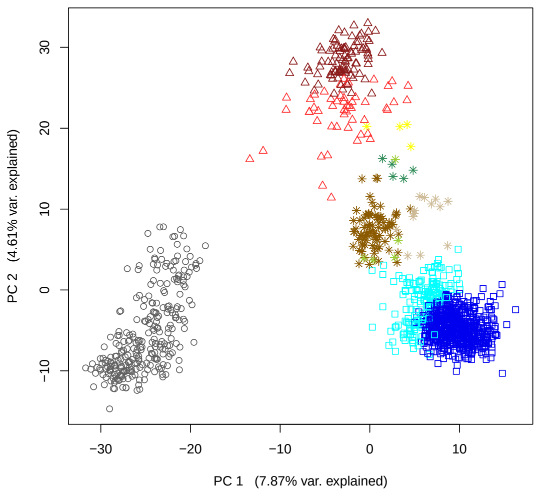


Figure S5: Principle Component Analysis of teosinte and Mexican maize samples. Colors and shapes correspond to taxonomic groups as follows: dark blue squares=high confidence parviglumis, light blue squares= ambiguous parviglumis, dark red triangles=high confidence mexicana, light red triangles=ambiguous mexicana, grey circles=maize, yellow pluses= central plateau hybrid group, brown pluses=central Balsas hybrid group, tan pluses= South Geurrero hybrid group, seagreen pluses= Huitzuco, and yellowgreen pluses=other hybrids

| | Parv_Balsas | Parv_Jalisco | Mex_Chalco | Mex_CP | Hyb_SG | Hyb_CB | Hyb_CP |
|---|---|---|---|---|---|---|---|
| Parv_Balsas | | 0.075 | 0.129 | 0.124 | 0.079 | 0.073 | 0.127 |
| Parv_Jalisco | | | 0.191 | 0.170 | 0.120 | 0.116 | 0.201 |
| Mex_Chalco | | | | 0.043 | 0.128 | 0.103 | 0.138 |
| Mex_CP | | | | | 0.097 | 0.086 | 0.091 |
| Hyb_SG | | | | | | 0.069 | 0.098 |
| Hyb_CB | | | | | | | 0.100 |
| Hyb_CP | | | | | | | |

Figure S6: All pairwise global $F_{ST}$s between hydrid groups, high confidence parviglumis, and high confidence mexicana

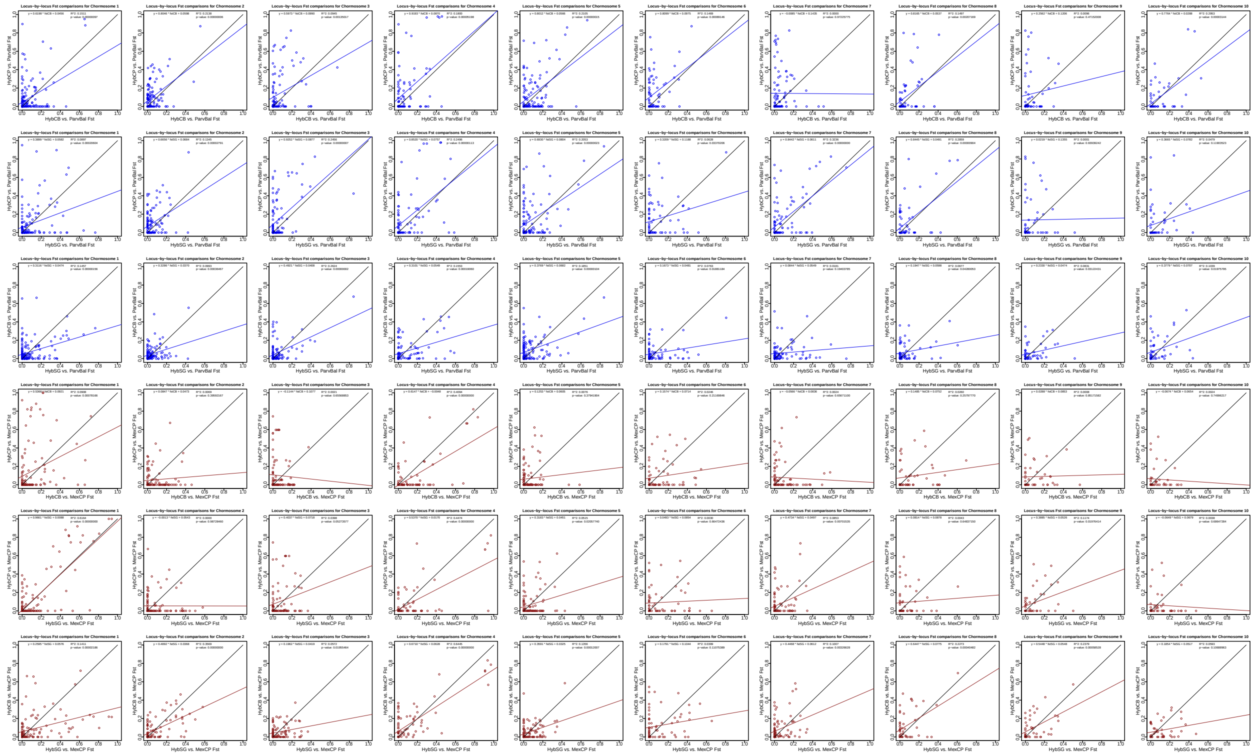

Figure S7: Comparisons of locus-by-locus $F_{ST}$ values between one hybrid group and another each compared to either parviglumis (blue) or mexicana (red) with one axis per hybrid group vs. mexicana for all ten chromosomes. A black line indicates y=x, and deviations from that line indicate a difference between the two hybrid groups with respect to a particular parental subspecies' alleles. A red or blue line indicates the linear fit line for all points. Also shown is the formula for the fit line, the $R^2$ value, and the p-value.

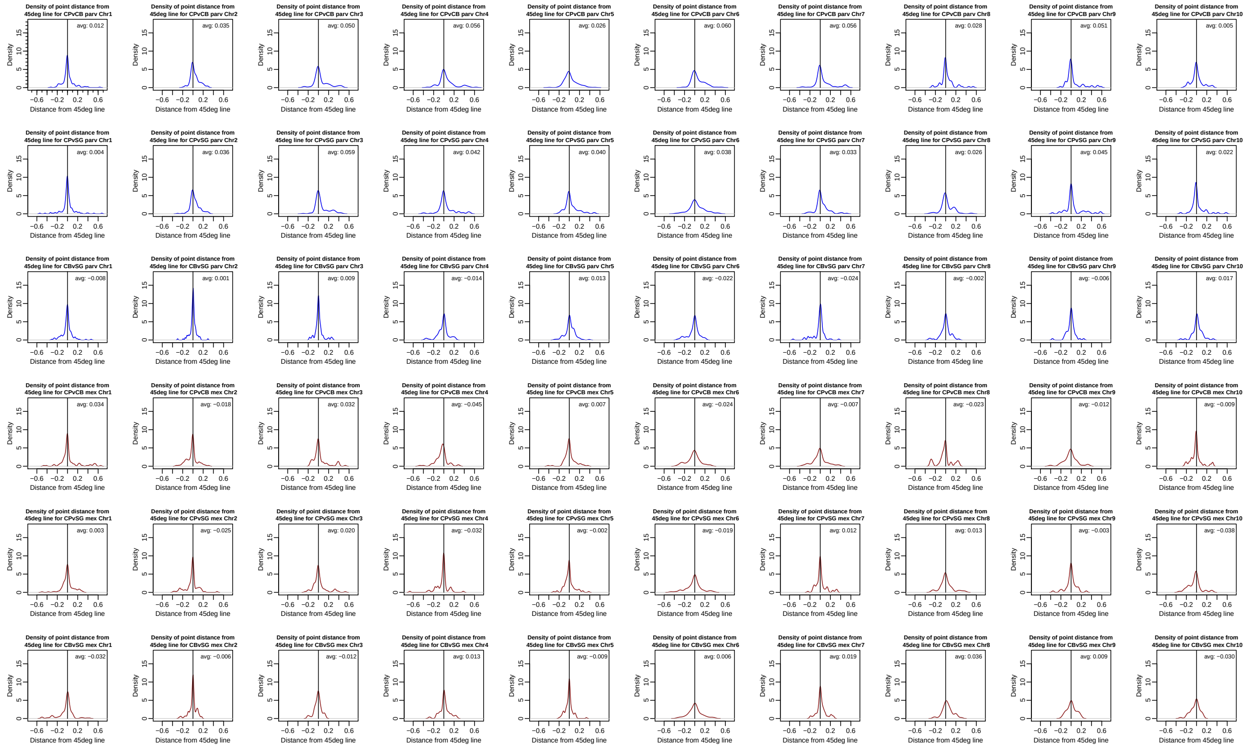

Figure S8: R density plot of deviations from the y=x line in Figure S7 indicating the difference between the two hybrid groups with respect to parviglumis (blue) or mexicana (red) alleles for all chromosomes. A black line shown is at zero to show the null expectation of each hybrid group having the same $F_{ST}$ compared to mexicana for all loci. Also shown is the averaged value from all data points to show the greater trend.

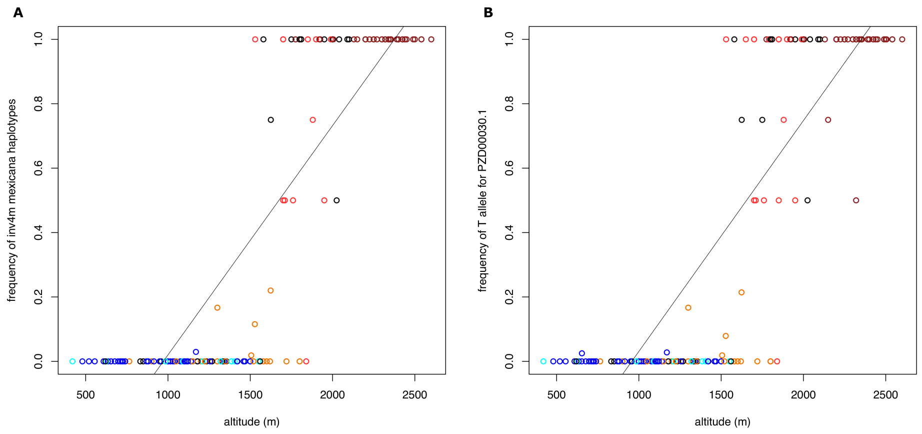


Figure S9: Frequency of A) inv4m mexicana haplotypes and B) PZD00030.1 T (mexicana) allele frequency over altitude. For both A and B: Each point represents a sampling site. Dark red indicates a population of all or mostly high confidence mexicana individuals, light red indicates ambiguous mexicana, dark blue indicates high confidence parviglumis, light blue indicates ambiguous parviglumis, orange indicates high confidence hybrids, and black indicates a mixed population. In order to be considered as having "mixed" hybrid status a sampling site must have at least 20% minority hybrid status individuals. Haplotype frequency was calculated by summing the haplotypes, for each sampling site, M weighted as two, P weighted as zero, and H weighted as one normalized by 2N where N is the sum of M, P and H in the sampling site. R is excluded from the calculation and therefore treated as missing data. PZD00030.1 is the highest $F_{ST}$ SNP on inv4m between high confidence parviglumis and high confidence mexicana

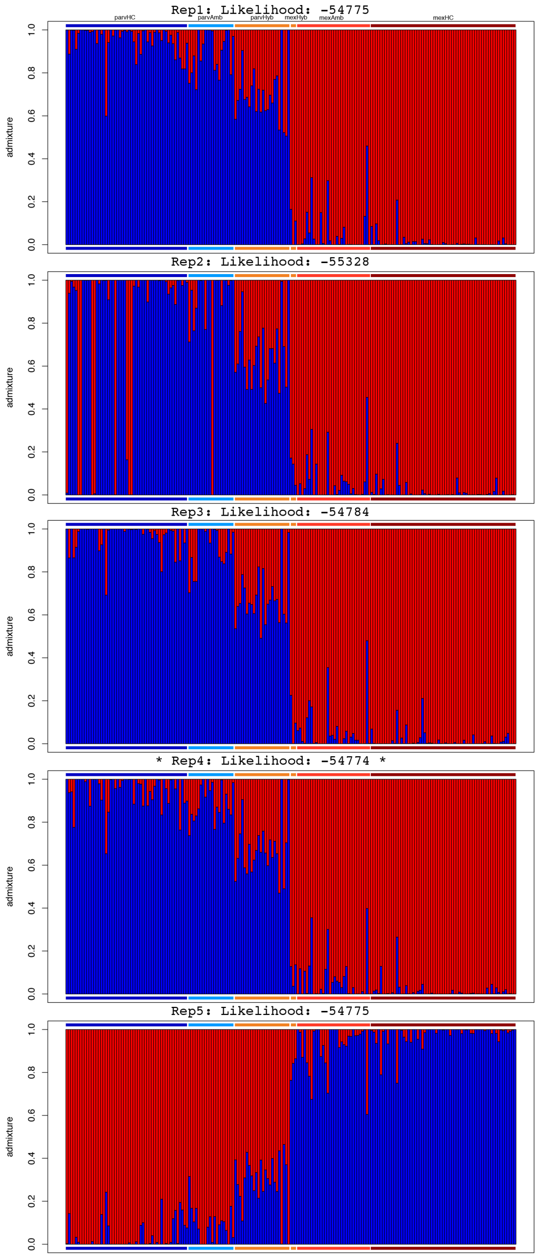


Figure S10: Five replicates of STRUCTURE-like bar plots generated using the spatial model of conStruct at group size two with model likelihoods above each bar plot. Each vertical line shows the attribution of an individual to parviglumis (blue) and to mexicana (red) based on per loci group assignments for all loci in the data set. Individuals within bar plots are ordered by their group as follows: high-confidence parviglumis, ambiguous parviglumis, South Guerrero hybrids, Central Balsas hybrids, ungrouped parviglumis hybrids, Central Plateau hybrids, ungrouped mexicana hybrids, ambiguous mexicana, and high-confidence mexicana. Colored horizontal bars above and below each plot represent these groups.
